# Is shed hair the most effective non-invasive resource for estimating wild pedigrees?

**DOI:** 10.1101/714964

**Authors:** Anubhab Khan, Kaushal Patel, Subhadeep Bhattacharjee, Sudarshan Sharma, Anup N Chugani, Karthikeyan Sivaraman, Vinayak Hosawad, Yogesh Kumar Sahu, Goddilla V Reddy, Uma Ramakrishnan

**Author notes:** Corresponding Author: Anubhab Khan, National Centre for Biological Sciences, TIFR, Bellary Road, Bangalore 560065, India.

## Abstract

Wild pedigrees are critical for better understanding mating systems and inbreeding scenarios to inform conservation strategies for endangered species. To delineate pedigrees in wild populations, many identified individuals will have to be genotyped at thousands of loci, mostly from non-invasive samples. This requires us to quantify (a) the most common non-invasive sample available from identified individuals (b) the ability to acquire genome-wide data from such samples, and (c) the quality of such genome-wide data, and its ability to reconstruct relationships between animals within a population. We followed identified individuals from a wild endangered tiger population, and found that shed hair samples were most common compared to fecal samples, carcasses and opportunistic invasive samples. DNA extraction, library preparation and whole genome sequencing resulted in between 126,129 and 512,689 SNPs from across the genome for four such samples. Exploratory population genetic analyses revealed that these data were free of holistic biases, and could recover expected population structure and relatedness. Mitochondrial genomes recovered matrilineages as suggested by long-term monitoring data. Even with these few samples, we were able to uncover the matrilineage for an individual with unknown ancestry. In summary, we demonstrated that non-invasive shed hair samples yielded adequate quality/quantity DNA AND in conjunction with sensitive library preparation methods, provided reliable data from hundreds of thousands of SNPs across the genome. This makes shed hair are an effective resource for studying individual-based genetics of elusive endangered species.

## Introduction

Long term monitoring of individuals and their relationships within populations provides key insights into demography, reproductive success, fitness and social organization (Pemberton, 2008; Kruuk & Hill, 2008). In particular, pedigree reconstruction in endangered populations is crucial for evaluation of inbreeding and mating patterns. Pedigree information from red wolf (*Canis rufus*) was used to quantify introgression with coyotes (Miller et al., 2003), while similar data in a wild wolf population (Liberg et al., 2005) quantified the effect of inbreeding on wolf pup survival in winter. Pedigrees have also been used to assess runs of homozygosity, or chromosomal stretches that might contribute to inbreeding depression (Kardos et al., 2018). Ongoing habitat fragmentation has resulted in small and isolated populations for many carnivores (Haddad et al., 2015; Crooks, 2002), and evaluation of pedigrees is becoming an increasingly important part of conservation planning and management, especially for large mammals.

Estimating pedigrees often requires tracking and following individuals, and monitoring their reproductive success. Molecular genetic data are essential to investigate paternity (Slate et al., 2000). Recent studies have also used genome-wide data to investigate paternity and population level pedigrees (Huisman, 2017; Hadfiled, 2012). Typically, such studies require wild individual capture and tagging and blood sample collection (Clutton-Brock & Pemberton., 2004). While this approach seems possible for some herbivores, it is difficult to implement for large elusive carnivores or endangered and rare species. In most cases, immobilization maybe additionally difficult or dangerous. For such species, minimally invasive samples like fecal matter (Solberg et al., 2006), excreted waste, pellets, saliva swabs from kill sites, environmental DNA or samples of shed skin, feather (Horvath et al., 2004), antler and hair are more feasible (Rozhnov et al., 2009). Unfortunately, most of these samples yield low quantities of DNA (Ball et al., 2007; Gupta et al., 2013). Such non-invasive samples are variably enriched in host DNA depending on the sample source. For example, whole DNA from scat samples are dominated by bacterial DNA (Chiou & Bergey, 2018) and prey DNA, while urine samples (if not already mixed with environmental DNA from soil or surface) have low amount of DNA and allelic dropout in microsatellite data (Carguilo et al., 2015). Saliva samples from kill sites might belong to more than one individual, as well as have bacterial and prey DNA contamination. Shed hair samples are expected to be enriched in host DNA but are potentially scarce at a site. Hair samples have been used to sequence and assemble whole genomes of extinct woolly mammoths (Miller et al., 2008). Here we attempt to identify the most common non-invasive samples in the field from identified individuals and the utility of these samples in recovering the pedigrees of wild populations. In order to do so, we sampled shed hair, scat, carcasses and blood (both opportunistically) from individuals in a wild tiger population.

Tigers are elusive and endangered large felids, making it difficult to sample them invasively. Because tigers have unique stripes, it is possible to identify individuals visually. First, we investigated the most frequently encountered samples from identified individuals and test whether these samples (a) can be collected optimally and (b) yield more genome-wide information than other non-invasive or invasive samples in the context of identified individuals. Finally, we assessed whether the genome-wide data generated provide biologically meaningful insights by investigating (a) documented/known patterns of population structure and (b) one case each of known and unknown maternity and relatedness.

## Materials & Methods

### Zoo and field sampling

Samples from a wild-caught tiger housed in a lone enclosure in a zoo was collected to optimize DNA extraction and sequencing. Shed hair in scratch marks on trees and on the ground where the tiger had been resting were collected. In the wild (Ranthambore Tiger Reserve), sampling was conducted as depicted in Box 1A. We sampled 38 wild tigers for 75 days (20^th^ May to 30^th^ June and in the month of November in 2017) and then again 180 days from 1^st^ January to 31^st^ June 2018. Fecal samples from individuals were collected by swabbing the surface of the scat using a sterile swab dipped in Longmire’s buffer (Longmire et al. 1997) and preserved in Longmire’s buffer until further processing.

**Box 1A.**
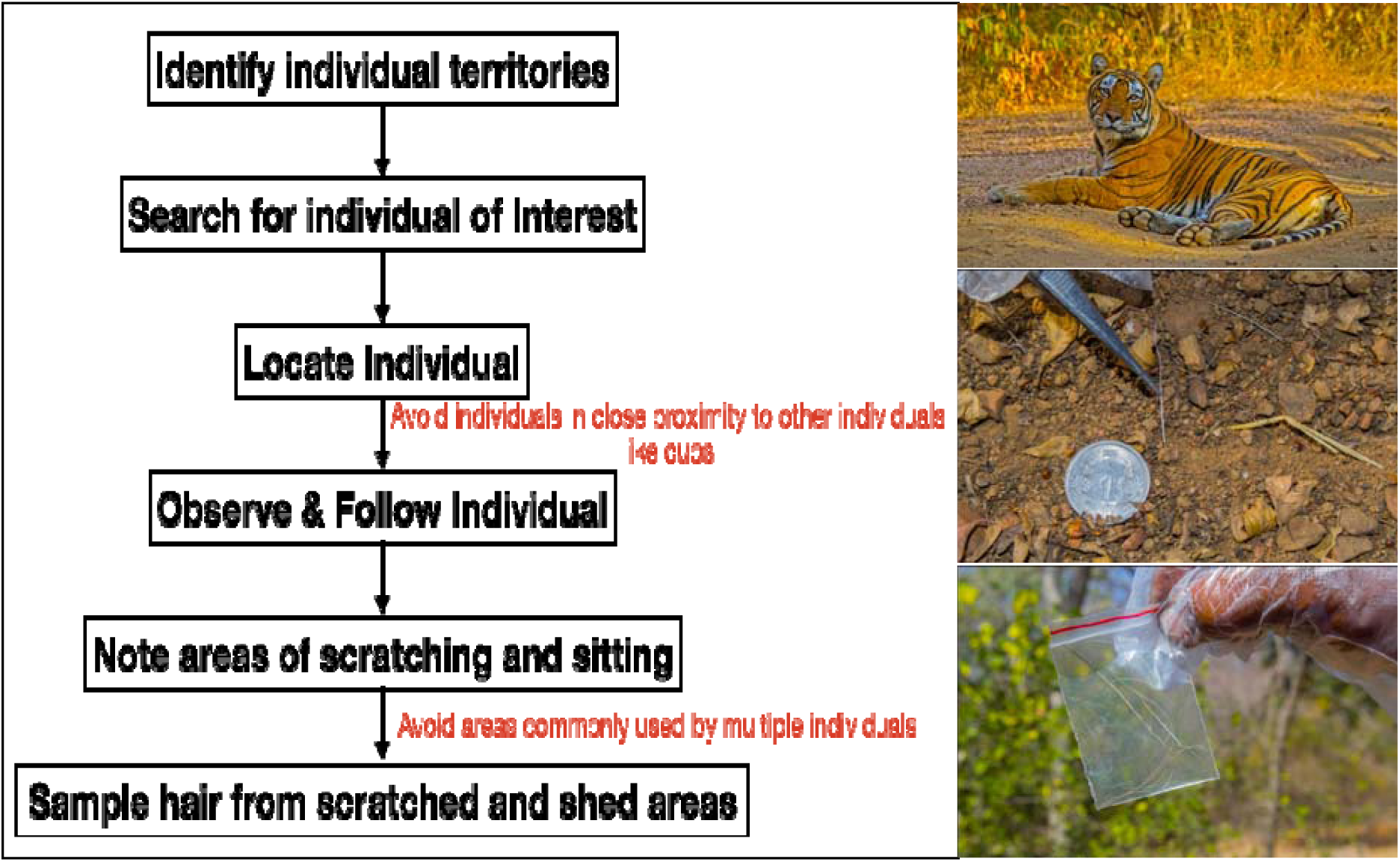
Hair sampling protocol used in this study. The text in red are the cautions to be followed in those steps. The diameter of the 1rupee coin is 2.193 cm

Tissues from the carcass of one of the individuals, T16, was collected in absolute ethanol and transported to the lab in gel packs.

### Laboratory methods

For the scope of this study, we used shed hair from 5 tigers; T24, T20, T47, T64 and T104; tissues from T16 and T104 and fecal samples from T03, T08 and T47. Supplementary table 1 lists the individuals and the kind of samples used in this analysis.

#### DNA extraction

The DNA was extracted based on the approaches depicted in Box 1B. Briefly, for the hair root only method, 10 hair roots were selected from the zoo individual and the hair shaft was discarded. To these, 200ul of AL buffer, 40ul of Proteinase K and 20 ul of 1M DTT were added and incubated overnight at 56°C. These hair roots were extracted using a modified protocol of the Qiagen blood and tissue extraction kit (Cat. 69504). DNA from hair root was extracted for tigers T24 and T47.

**Box 1B.**
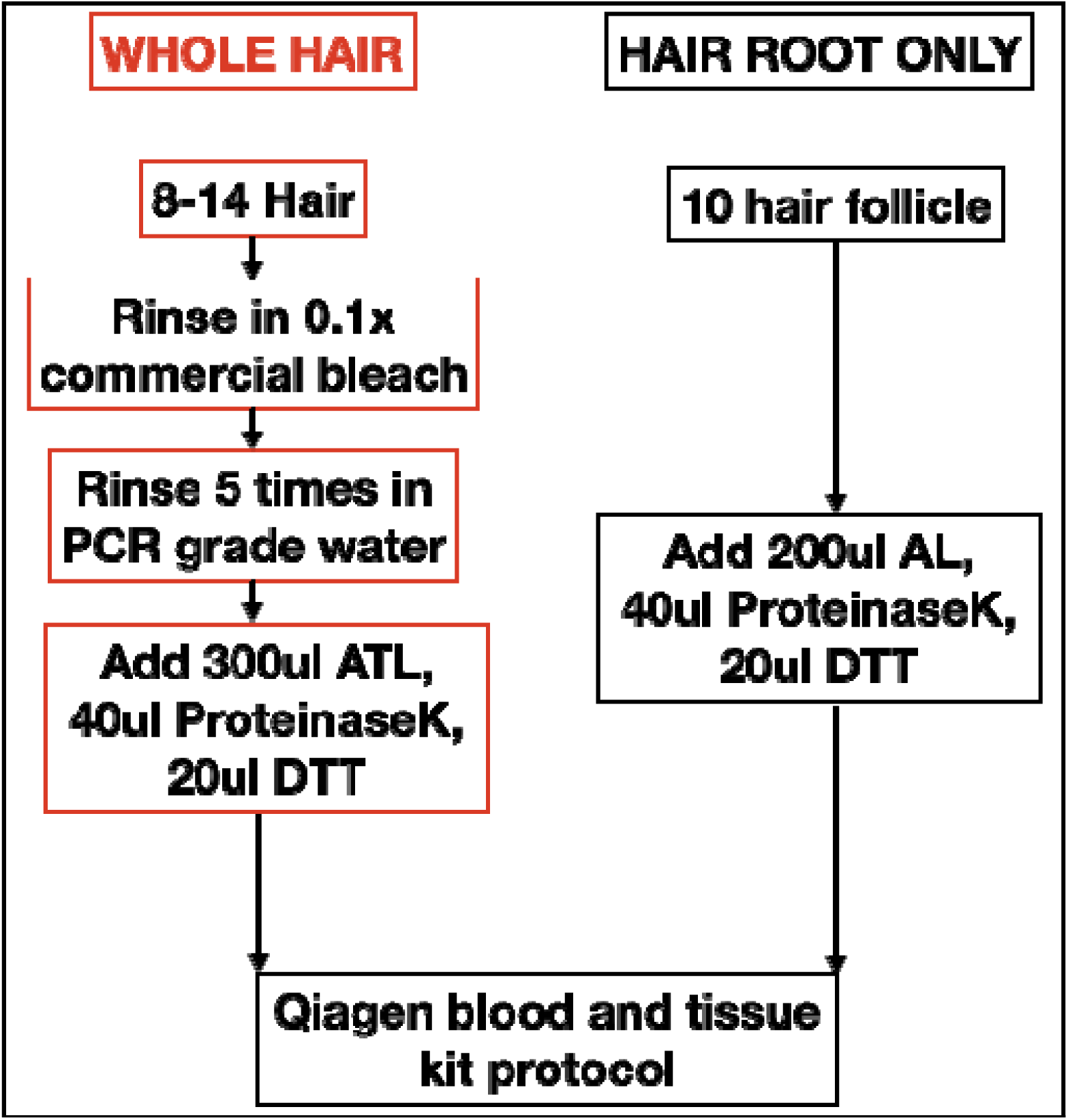
DNA extraction protocols used for whole hair and hair root.

For the whole hair DNA extraction, 8-14 randomly selected hair strands were rinsed in 0.1X commercial bleach and washed with nuclease free water. To this, 300 ul ATL buffer (Qiagen), 30ul ProteinaseK, 20 ul 1M DTT was added and incubated at 56°C until visible lysis. To this lysate 300ul AL buffer (QIAGEN), 3ul of 1ug/ul carrier RNA, 300ul of absolute alcohol was added (in that order), vortexed and loaded onto the spin column. The rest remains the same as mentioned in Qiagen blood and tissue extraction kit handbook. DNA from whole hair was extracted for tigers T20, T24, T47, T64 and T104.

DNA from tissue samples of tigers T16 and T104 was extracted using Qiagen blood and tissue extraction kit (Cat. 69504) as per the manufacturer’s instructions. Fecal samples from T47 and T03 were extracted using the method described in Natesh et al. (2019).

#### Library preparation and Sequencing

DNA whole genome libraries were prepared using NEBNext® UltraTM II DNA Library Prep Kit (#E7645L, NEB Inc). DNA was quantified on QubitTM 3.0 flourometer using Qubit High sensitivity dsDNA Assay (#Q32854, Thermo Fisher Scientific). The quantified libraries were then clonally amplified on a cBOT and sequenced on the HiSeq X with 150bp paired end chemistry.

### Analyses

The reads from 150 bp paired end sequencing were trimmed using TRIMMOMATIC (Bolger, Lohse & Usadel, 2014) to have an average PHRED quality of 30 in a sliding window of 15 bp, and any read that was shorter than 36 bp after trimming was removed from further analysis. These reads were then aligned to 1) the tiger genome assembly (Armstrong et al., 2019) and 2) the mitochondrial genome of tiger (NC_010642.1) using BOWTIE2 (Langmead & Salzberg, 2012). The alignments were then saved in a binary format using SAMTOOLS (Li et al., 2009). To assess the quality of the alignments QUALIMAP (Garcia-Alcalde et al., 2012) was used.

### Sample dependent data quality

To test difference in data quality across different kinds of samples, we used data from whole genomes from the tissues used here, whole genomes from shed hair and whole genomes sequenced from fecal extracts (for tigers T03 and T47). Data from samples with a higher number of reads was subsampled to match the samples with the lowest number of reads. Comparison of the raw data without controlling for number of reads is presented in Supplementary table 2.

To test the differences in sequences obtained due to use of different samples, we estimated the percent number of sites out of the called SNP sites that were called erroneously. We estimated mismatches (0: identical; 1: single allele mismatch; and 2: both alleles mismatch) between fecal genome SNPs, hair root genome SNPs and whole hair genome SNPs for tiger T47. In one other case, we compared SNPs called from whole hair and blood from tiger T104. We subsampled the vcf files to contain only the samples being compared (Supplementary table 3). For fecal vs hair root genome and fecal versus whole hair genome we obtained 1,213,803 SNP loci with no missing data. For hair root genome versus whole hair genome, we obtained 2,917,519 SNP loci and for whole hair genome versus blood genome we obtained 4,353,417 SNP loci with no missing data. On these files we used the –genome function of PLINK (Purcell et al. 2007) to obtain the pairwise mismatches.

### Sample dependent data bias

To test for biases in the sequencing, ddRAD data generated earlier (Natesh et al., 2017) for three tiger reserves, namely, Kanha Tiger Reserve, Wayanad Wildlife Sanctuary and Ranthambore Tiger Reserve was used. All reads were trimmed and aligned to the tiger genome as per the methods described here previously. Variants were called for the entire dataset of using bcftools multiallelic caller (Li et al., 2009). The raw variants were filtered for Quality and Genotype quality of 30 (this ensured we have 99.9% confidence in the bases and the genotypes), depth of 10, maximum missing data allowed per locus of 20%, conformity of hardy Weinberg equilibrium at a P value of 0.05 (this removed the improbable genotypes), minor allele count of 3 (this ensures that any alleles is present is at least 2 individuals, this removes any sequencing error) and also indels were removed (this ensured only SNPs were being used). To recover the population structure obtained by Natesh et al. (2017), we used the filtered SNP dataset. Structure was estimated using the programme fastSTRUCTURE (Raj, Stephens and Pritchard, 2014) for complexity values of 2,3,4 and 5. The complexity value with the maximum likelihood was chosen.

### Matrilineage analysis

We attempted to estimate the matrilineage of four tigers in our dataset. Behavioural observations from multiple independent sources and aging data on tigers (Sadhu et al., 2018) reveals several mother-offspring relationships and maternal sibships. Consensus mitogenomes from whole genome sequences were called using the Amur tiger mitogenome as a reference (RefSeq NC_010642.1). Multiple sequence alignments (MSA) of whole mitogenomes from T16, T20 and T64 were performed using clustal-omega (Sievers et al., 2011) and then a median joining network was created using popart (Leigh, Jessica and Bryant, 2015). The mitogenome sequence obtained using the shed hair genome was tested using the known matrilineage of T16, T20 and T64 while previously unknown matrilineage was inferred for T24, T47 and T104.

## Results

### We detail results in the following sections

#### Is shed hair an abundant and effective source of DNA and genome-wide data?

We followed 34 individual wild tigers identified from their unique stripe patterns (Supplementary Figure 1), and obtained shed hair samples from 207 sitting sites. 10 scats samples could be obtained from 9 of these individuals. Additionally, tissues from 3 tiger carcasses (death due to conflict) and one opportunistic tranquilization (Figure 1a) were obtained. From the 207 hair collection sites, we obtained on an average 25 hair strands per site (Figure 1b) of which approximately 65% of the hair strands had a potential hair root (Figure 1b).

**Figure 1.**
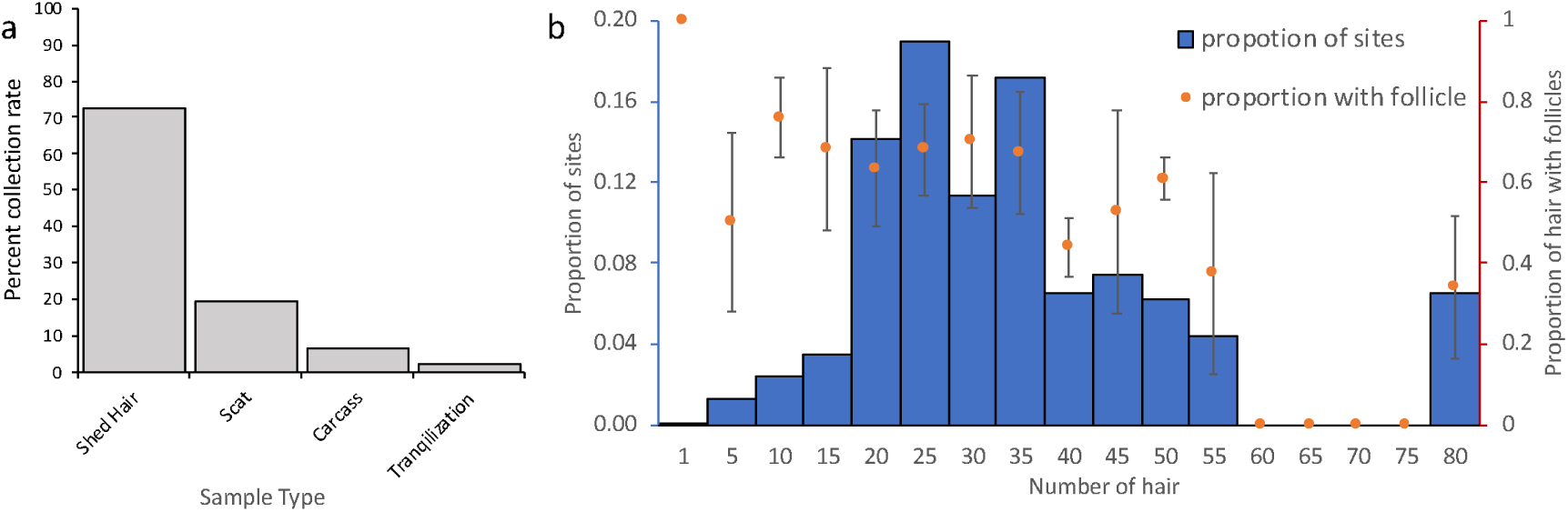
(a) In a period of 255 days of following 34 individuals, shed hair samples could be obtained for all 34 individuals while scat could be obtained for only 9. Opportunistically, 3 carcasses were recovered and 1 tranquilization was reported. (b) Number of hair strands collected per site. Out of the total hair strands collected per site, most sites seem to have at least 50% of the hair with follicle.

To evaluate the best strategy for DNA extraction and sequencing, shed hair from a wild caught tiger T24 housed in zoo and a wild tiger T47 were sampled. Sequencing of the DNA from hair root extracts yielded 13,452,410 and 373,791,866 reads while that from the whole hair yielded 15,735,782 and 341,232,300 reads (after adapter trimming) from T24 and T47 respectively. The whole hair DNA extract had more tiger DNA and less bacterial DNA (Supplementary Figure 2). The DNA from the whole hair had more percent mapped reads to nuclear and mitochondrial DNA of tiger and covered more of the genome. However, the duplication rate for reads aligned to nuclear and mitochondrial genome (indicating PCR duplicates) was higher in whole hair DNA extract (Figure 2A and 2B).

**Figure 2.**
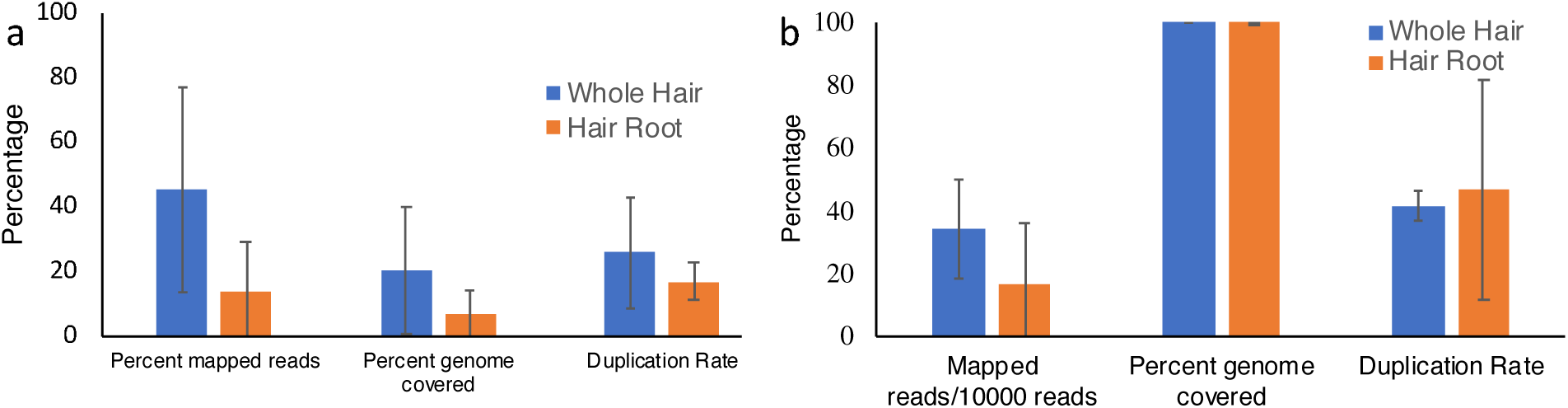
Whole hair DNA extracts have more reads mapping to the reference tiger (a) nuclear genome and cover more bases on the nuclear genome and also on the (b) mitochondrial genome. The number of reads obtained from all sequences have been normalized.

Across the five whole genomes of shed hair samples, the sequence quality was variable (Supplementary Table 2). The minimum percent nuclear genome covered in our dataset was 24.85 (yielding 126,129 SNP loci) for shed hair from the tiger T20 and was maximally 98.03 (yielding 512,689 SNP loci) for tiger T47. However, increasing the sequencing depth increased the percent genome covered.

To test how shed hair performed in comparison to DNA from tissue and scat, we compared DNA sequences from tissues (WGS data of tigers T16 and T104), whole genome sequences from shed hair and scat DNA ddRAD (ENA accession number ERP014988) combined with scat DNA WGS (tigers T03 and T47). The number of nucleotides sampled were normalized across all. Tissue samples performed best (in terms of mapped reads, and percentage genome covered), while scat samples performed the worst (Figure 3). The variance in the shed hair whole genome sequencing data was very high but overall, the average was better than scat.

**Figure 3.**
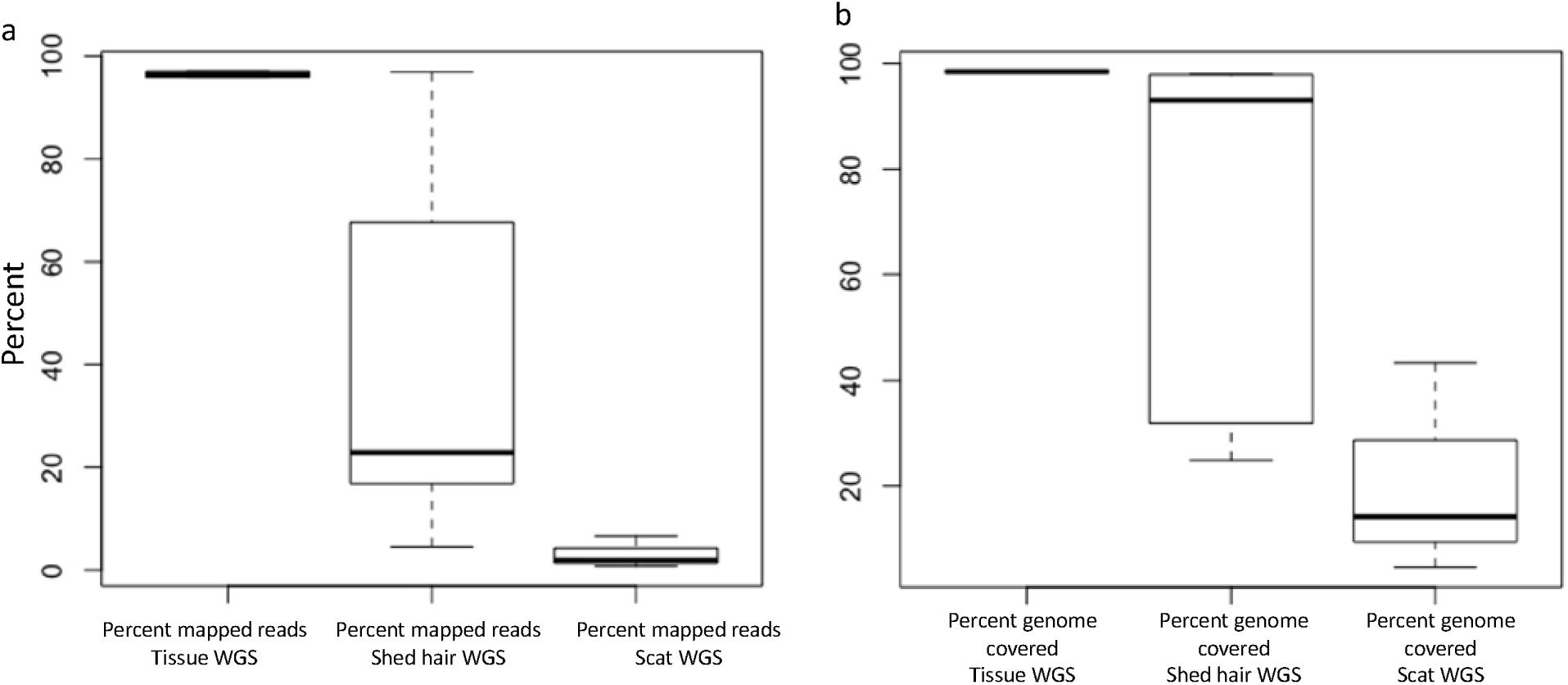
(a) Sequencing reads from tissue samples (n=2) have higher mapped reads and low variance to the tiger nuclear genome as compared to hair (n=5) or scat samples (n=3). Hair samples have very high variance. (b) Tissue samples perform better even in terms of percent genome covered even though the library preparation is a reduced representation of the genome.

#### Do genome-wide data from shed hair provide meaningful results?

We compared mismatches between different sample sources and whole hair versus blood comparisons revealed (for tiger T104) that 96% of the loci had no mismatches while 4% of the loci had a single mismatch (Figure 4). Scat versus hair root and scat vs whole hair (for tiger T47) had 47% and 37% of the loci respectively without mismatches, while 19% and 24% of the loci had 1 mismatch. Comparison between hair root and whole hair (for tiger T47) revealed that 64% of the loci had no mismatches while 29% of the loci mismatched for 1 allele.

**Figure 4.**
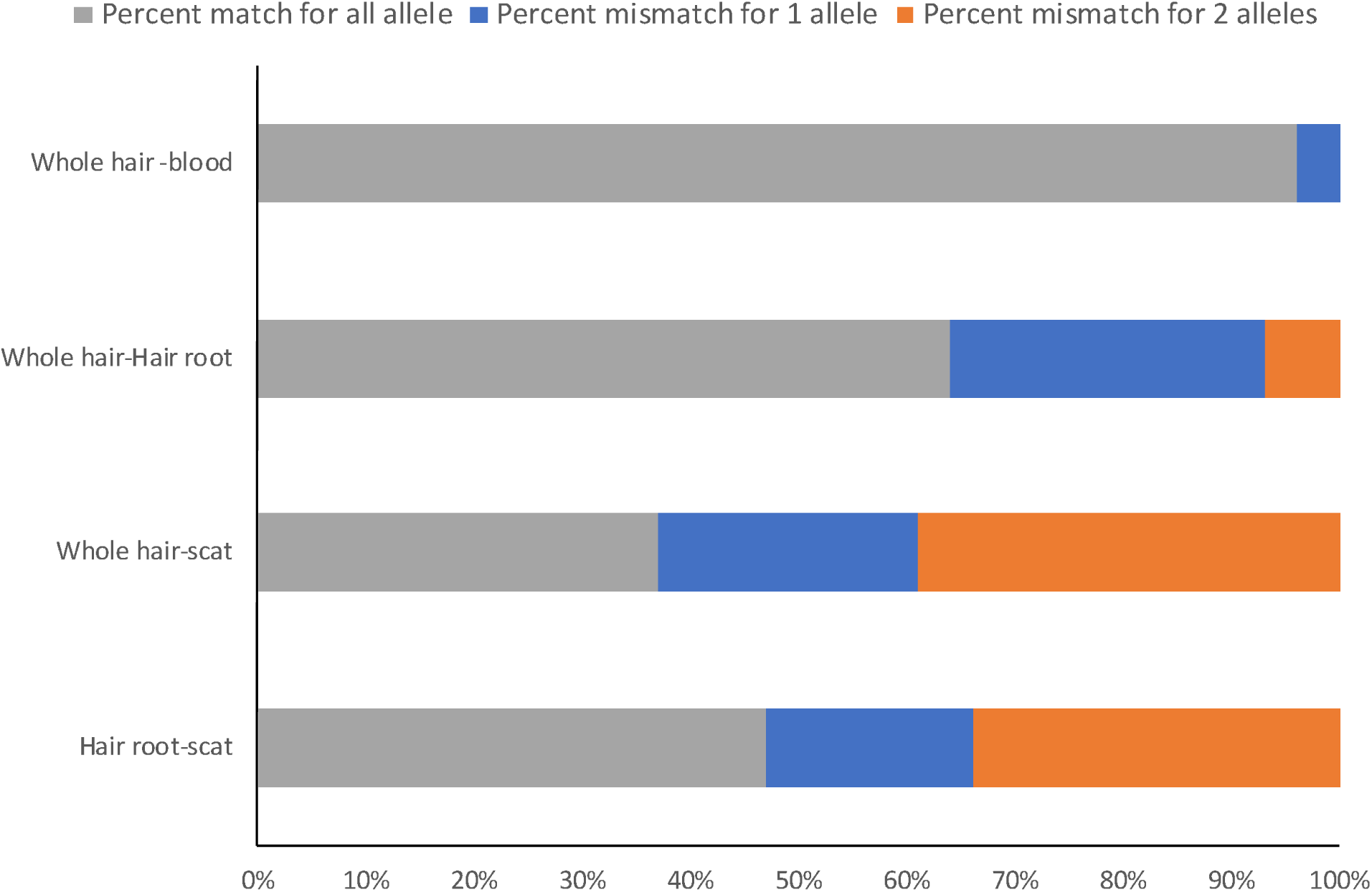
Percent pairwise mismatch between SNP data from different sample types.

We combined our data with those from three tiger populations identified in Natesh et al. (2017) viz. Kanha Tiger Reserve, Wayanad Wildlife Sanctuary and Ranthambore Tiger Reserve and obtained 15,644 SNPs from this combined dataset. fastStructure results replicated the optimum complexity of 3 (Figure 5a). We did not find any grouping (between sample types or otherwise) within Ranthambore (Supplementary figure 3). The relatedness estimates also revealed similar patterns, with Ranthambore having the highest average pairwise relatedness (Figure 5b).

**Figure 5.**
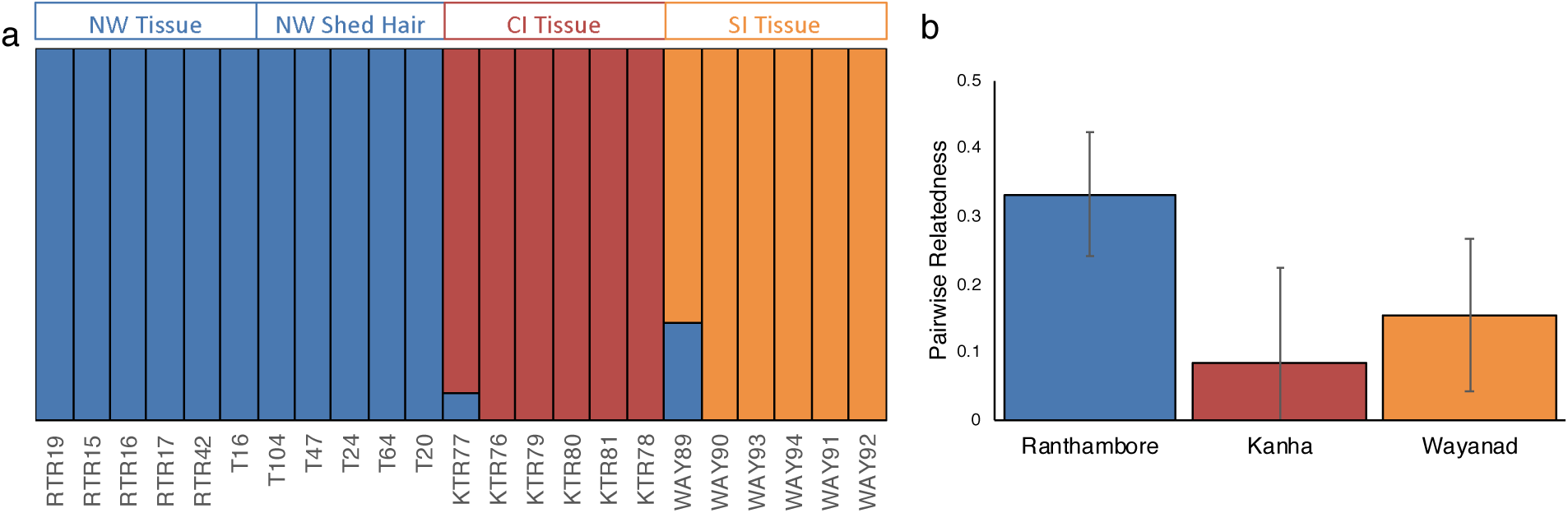
Results from Natesh et al. (2017) could be replicated after adding in the shed hair whole genome sequences. (a) If there were specific biases shed hair WGS samples would have formed a separate cluster. The optimal complexity was 3. SI, NW and CI stand for Wayanad Wildlife Sanctuary, Ranthambore Tiger Reserve and Kanha Tiger Reserve. (b) The trends in pairwise relatedness for tiger reserve is similar in our dataset and that in Natesh et al. (2017). Ranthambore has the highest pairwise relatedness among the tiger reserves here.

The tiger population in Ranthambore Tiger Reserve has been monitored closely by the forest department staff. Due to their efforts, maternity and sib-ship relationships are estimated for several tigers. From this the matrilineage of tigress T16 is thought to be one of the founders, and most tigers are supposed to have descended from her, making the relatedness between individuals high. However, certain tigers have no known maternity or matrilineage, making them an ideal test case. The tigers T20 and T64 are supposed to share T16’s matrilineage. While the mother of T24 is thought to be T22 and that of T104 is thought to beT41. T47’s maternity and matrilineage both are unknown.

As expected from the long-term data depicted in figure 6a, we find a linear network (ordered by generation), starting with T16 (figure 6b). Thus, we recovered known matrilineage reliably. Using data from all 6 individuals analyzed here, we obtained the haplotype network depicted in Figure 6c. The network suggests that T47 belongs to same matrilineage as T16, T20 and T64 while T24 and T104 potentially belongs to a different matrilineage. Additionally, pairwise relatedness (using 15644 SNPs from the nuclear genome) between T24 with others and T104 with others is lower than pairwise relatedness estimates between half/full siblings T20, T47 and T64. The maximum relatedness of T24 is to T16 at 0.35 while the minimum relatedness is of T104 with T24 at 0.25 and the maximum is for T20 -T47 pair at 0.65 (Figure 6d). Thus, T47 might be a previously unknown full sibling of T20 and both sons of T16.

**Figure 6.**
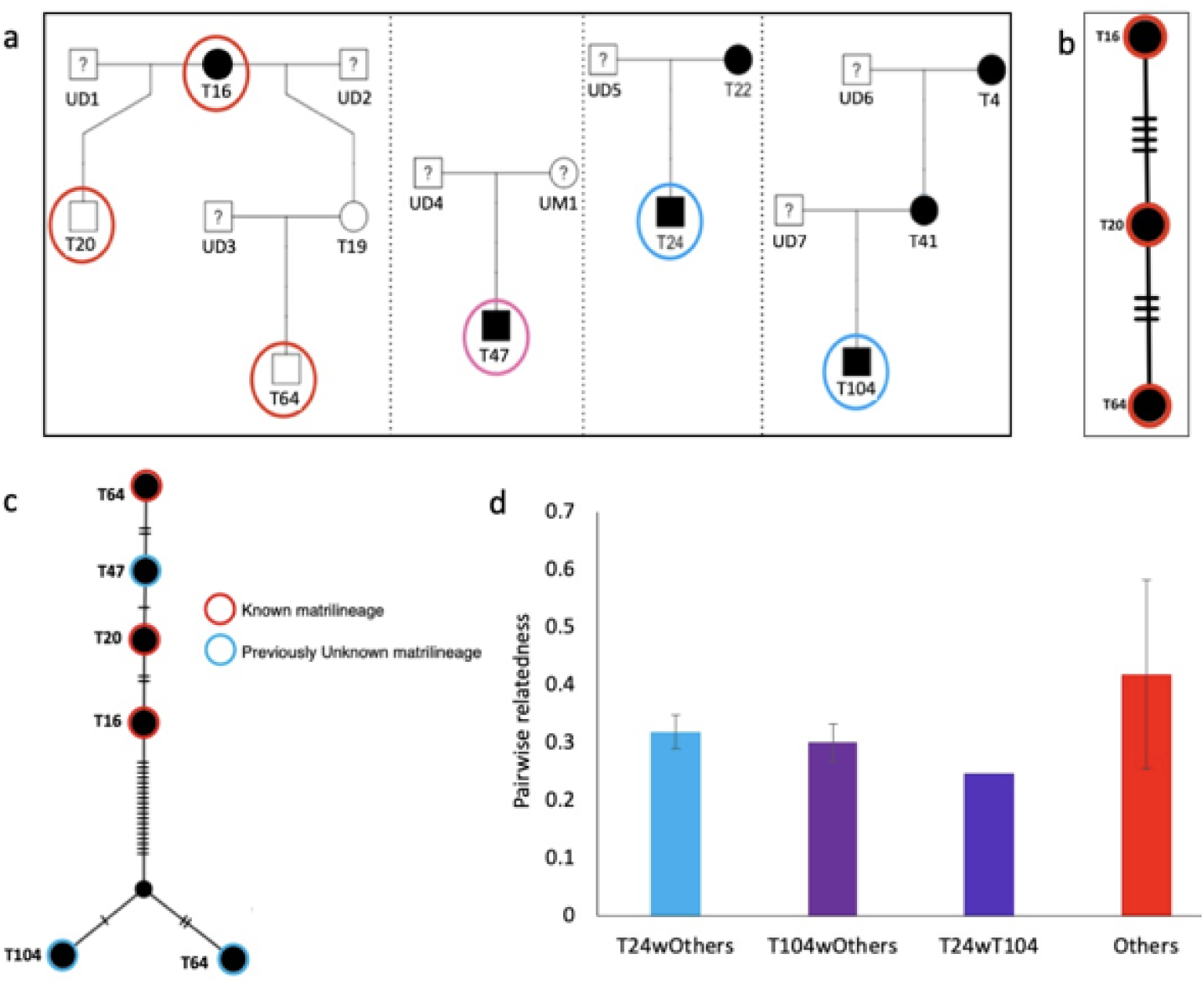
(a) The estimated pedigree of RTR tigers from behavioural observations. The individuals in circles are sampled here and the individuals in red samples have known ancestry. (b) The estimated minimum spanning network for mitochondrial genome for T16, T20 and T64. This is consistent with the behavioural data. (c) The estimated minimum spanning network for mitochondrial genome for all individuals in our dataset. (d) The pairwise relatedness using nuclear SNPs.

## Discussion

Here we establish that shed hair from identified individuals is an adequately available and effective source of DNA for generating genome-wide data and estimating pedigrees. Thus far, such individual-based molecular studies have been conducted mostly with captured and tagged individuals involving invasive sampling (e.g. Soay sheep, red deer (Clutton-Brock & Pemberton, 2004), meerkats (Leclaire et al., 2013; Ross-Gillespie & Griffin, 2007) and Wolves (Vonholdt et al., 2008)) or with behaviourally invasive baited hair traps (e.g. red fox, Vine et al., 2009; black bear, Gardner et al., 2010; marten, Mowat and Paetkau, 2002; Eurasian lynx, Davoli et al., 2013; Southern hairy-nosed wombats, Walker et al., 2008 and Ocelots, Weaver et al., 2005). Our results reveal that shed hair is a viable sample source for individual-level genetic studies. Shed hair has rarely been used (e.g. captive Panda, *Ailuropoda melanoleuca* (Durnin et al., 2007) and wild Orang-Utans, *Pongo pygmaeus*, (Goossens, Abdullah, Sinyor, 2004)). We show that shed hair is a “high throughput” non-invasive sample compared to scats from identified individuals or carcasses. We suggest that collection of shed hair may allow individual and population level whole genome based studies in a shorter span of time. This is especially important for conservation biology studies since scientifically informed decisions are often delayed due to difficulties in collecting identified individual samples.

The population described here is one of the few high tiger density populations (e.g. Karanth et al., 2004). This contributes to the sampling rates we report here. For populations with low carnivore densities or difficult terrains, baited camera traps might be a better option for collecting hair. Hair traps in conjunction with camera traps can be used to collect samples from identified individuals especially in the case of species with non-pelagic patterns. Similarly, individual level sampling rates are also expected to be variable and in some cases, baiting may help.

Ecological and evolutionary genetics studies can benefit greatly from advances in next generation sequencing methods. However, obtaining good samples for wild individuals has always been a challenge. Using good non-invasive samples can solve a lot of problems. Methods that allow non-invasive samples to be used for obtaining genomic scale data like host DNA enrichment in fecal samples (Chiou and Bergey 2018), salivary samples from predatory bite marks (Blejwas et al. 2006) or baited camera traps (Shardlow and Hyatt 2013) will aid the most. For optimal usage of shed hairs, better methods of DNA extraction are needed. The host DNA yield from 10-12 shed hair is often low and cannot be quantified easily. Hair metagenome is known to have several contaminating DNA and more so in the hair roots owing to its relatively porous nature (Miller et al., 2008). Methods that can increase the efficiency of lysis in conjugation with enrichment methods will reduce contamination, thus increasing the overall host DNA content. This will have a twofold advantage of reducing potential sources of bias during the analysis and yielding more usable sequence data per unit raw data. The Chelex extraction method used by Bjornerfeldt and Vila (2007) used to obtain DNA from single hair needs to be tested on shed hair too but it would be significantly more expensive than the method described here. Advances in low DNA concentration library preparation method followed by short read sequencing will enable one to use non-invasive samples more effectively. Whole genome sequences from non-invasive samples will help in accurate and faster studies quantifying inbreeding using runs of homozygosity, identifying adaptive or deleterious alleles, identifying functional genomic regions for endangered charismatic mammals.

Our results point to at least one new matrilineage in Ranthambore Tiger Reserve (RTR). This population has undergone several bottlenecks most recent one in the year 2005, with few associate founders including T16. Bottlenecks are known to reduce allelic diversity and hence one might expect lower numbers of mitochondrial haplotypes in RTR compared to other population. Singh et al. (2013) have studied the dispersal of tigers in the landscape however, no evidence has so far been presented on immigration of tigers into RTR. In such a scenario discovering a previously unknown matrilineage of tigers in RTR suggests the potential for additional founder lineages or maybe, hitherto undocumented dispersal. The matrilineage of the tiger T24 was inherited from his mother T22 whose presence was detected as a prime adult in 2006 (Sadhu et al., 2017) and its whereabouts before this are unknown. Similar is the case for tiger T4 who is the grandmother of T104. Such lineages can supplement the population with genetic variation. Recovering matrilineages can prove to be important when estimating the pedigree of a population using several SNP markers from the nuclear genome. The matrilineages can be used as priors for the estimation of pedigrees else they can be used for annotating the pedigree recovered from SNP markers. Thus discovering matrilineages undocumented in observation based data is important for recovering a wild pedigree.

Although we added samples to the Ranthambore dataset from Natesh et al., (2017) the pairwise relatedness based on several thousands of SNPs between individuals within this population remains high. This suggests the possibility of inbreeding in this population, and that most individuals are highly related. Actual inbreeding can be tested using methods to estimate runs of homozygosity, based on samples collected from several individuals as described here. Inbreeding and inbreeding depression (if any) needs to be estimated and incorporated into management plans for this and other such isolated populations. The same sampling strategy could allow us to estimate the pedigree using genotypes of several individuals, and correlates of inbreeding depression by estimating differences in the number of successful offspring between inbred and outbred individuals. Such studies will allow us to investigate inbreeding avoidance, heritability of traits, heritability of territories, correlation of life history traits and genotype and several others for tigers.

## Conclusions

We aimed to discover the best sample types for studying genetics of identified individuals and if such samples can actually be used for genetic studies. We find that shed hair samples from identified individuals are the most available sample types and DNA sequences from whole shed hair is better than using only the hair roots. We establish that the sequences obtained from whole hair are reliable and match 96% of the sequence obtained from blood. However, we do find large variations in the amount of data obtained from whole shed hair, and that the DNA obtained is generally low concentration. In the future, it might be possible to also use probe-based approaches to extract information on specific loci and/or genomic regions to enable most appropriate use of she hair samples. In summary, we suggest that shed hair from identified individuals is a viable source of genome-wide data at the individual level.

## Acknowledgements

This work was supported by the Wellcome Trust/DBT India Alliance Senior Fellowship [grant number IA/S/16/2/502714] awarded to Uma Ramakrishnan. Permissions for sample collection were granted in letter number 19 (Part-Uma) Permission/Research/CWLW/2017 date 15/12/2017/300. Mr. Girish Panjabi, field ecologist, Ranthambore Tiger Reserve, helped us in identifying individuals. Gratitude to field assistants and several volunteers for their continuous presence in the field and assisting in sample collection for this project.

