## Supplementary information for "Is shed hair the most effective non-invasive resource for estimating wild pedigrees?"

**Supplementary tables and figures**

**Supplementary table 1:** The samples used in present study

| Individuals | Samples collected | DNA sequence source |
| --- | --- | --- |
| T16 | Muscle Tissue from corpse | Tissue |
| T20 | Shed hair | Whole Hair |
| T64 | Shed hair | Whole Hair |
| T24 | Shed hair | Whole Hair |
|  |  | Hair Root |
| T47 | Shed hair | Whole Hair |
|  |  | Hair Root |
|  | Scat | Fecal |
| T03 | Scat | Fecal |
| T08 | Scat | Fecal |
| T104 | Shed hair | Whole Hair |
|  | Blood from tranquilization | Tissue |

**Supplementary table 2:** Sample sequencing data quality

| Individual | DNA source | Percent mapped reads | Percent genome covered | Total number of reads |
| --- | --- | --- | --- | --- |
| T104 | Whole hair | 96.91 | 97.91 | 332,479,846 |
| T20 | Whole hair | 4.51 | 24.85 | 162,272,498 |
| T64 | Whole hair | 16.79 | 31.89 | 343,777,966 |
| T47 | Whole hair | 67.68 | 98.03 | 341,232,300 |
| T24 | Whole hair | 22.87 | 93.04 | 393,167,528 |
| T47 | Scat | 6.63 | 43.32 | 185,253,494 |
| T03 | Scat | 1.94 | 14.2 | 177,066,138 |
| T8 | Scat | 0.77 | 4.56 | 132,602,774 |
| T104 | Tissue | 97.05 | 98.49 | 304,471,110 |
| T16 | Tissue | 95.89 | 98.49 | 710,167,210 |

**Supplementary table 3:** SNP filters and individuals used in specific analysis

| Type of Analysis with DNA sequence data | Tool | VCF filters | Samples used | Number of SNPs |
| --- | --- | --- | --- | --- |
| Percent Mismatch of scat with whole hair | PLINK | MinQ 30, minGQ 30, minDP 10, mm 0, rmvIndels | T47 WH, T47 SC | 1,213,803 |
| Percent Mismatch of scat with hair root | PLINK | MinQ 30, minGQ 30, minDP 10, mm 0, rmvIndels | T47 SC, T47 HR | 1,213,803 |
| Percent mismatch of whole hair with hair root | PLINK | MinQ 30, minGQ 30, minDP 10, mm 0, rmvIndels | T47 WH, T47 HR | 2,917,519 |
| Percent mismatch of whole hair with tissue | PLINK | MinQ 30, minGQ 30, minDP 10, mm 0, rmvIndels | T104 WH, T104 TS | 4,353,417 |
| Structure between whole hair and tissue | fastSTRUCTURE | MinQ 30, minGQ 30, minDP 10, mm 0.8, hwe 0.05, mac 3, rmvIndels | Data from Natesh et al 2017, T16 TS, T104 WH, T47 WH, T64 WH, T20 WH, T24 WH | 15,644 |
| Within Ranthambore structure between samples | fastSTRUCTURE | MinQ 30, minGQ 30, minDP 10, mm 0.8, hwe 0.05, mac 3, rmvIndels | RTR Data from Natesh et al 2017, T16 TS, T104 WH, T47 WH, T64 WH, T20 WH, T24 WH | 15,645 |
| Pairwise relatedness | PLINK | MinQ 30, minGQ 30, minDP 10, mm 0.8, hwe 0.05, mac 3, rmvIndels | Data from Natesh et al 2017, T16 TS, T104 WH, T47 WH, T64 WH, T20 WH, T24 WH | 15,646 |

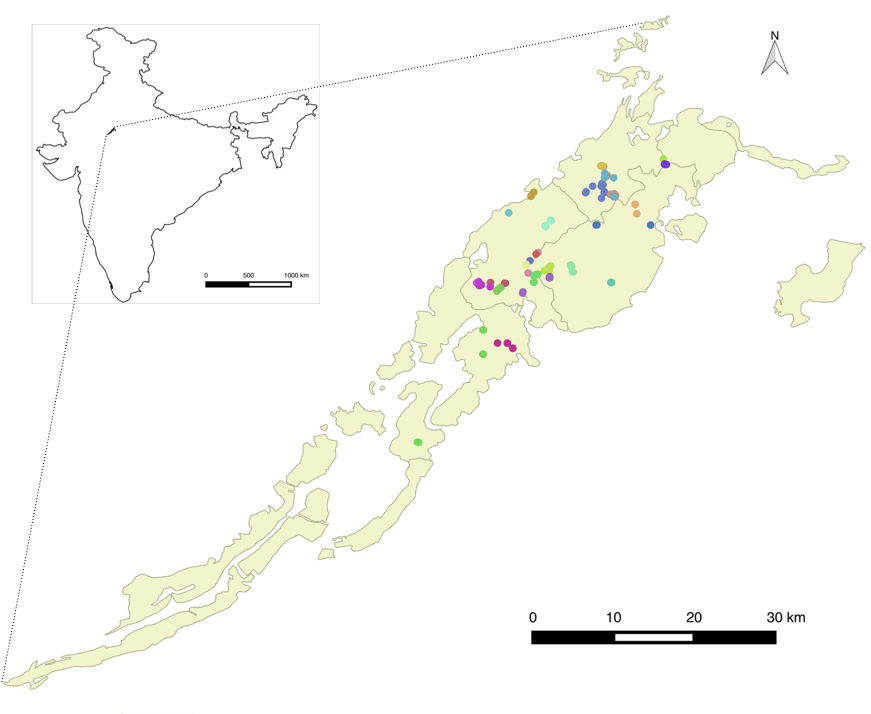

**Supplementary figure 1:** Study location. Ranthambore Tiger Reserve is in north-western India. Every dot on the map represents a sampling location and different colours represent different individuals. A total of 34 individuals were sampled.

Percent

| samples | Percent mapping to tiger genome | Percent mapping to human genome | Percent mapping to mouse genome | Percent mapping to Bacteria |
| --- | --- | --- | --- | --- |
| T20 | 3.94 | 32.68 | 3.7 | 3.78 |
| T24_whole_hair | 22.27 | 44.7569 | 0.00659461 | 1.02196 |
| T24_follicle_only | 2.41 | 10.6373 | 0.00869527 | 3.17346 |
| T47 | 60.8 | 9.52168 | 0.100906 | 0.269154 |
| T64 | 19.13 | 50.2122 | 0.649946 | 0.741194 |

**Supplementary figure 2:** Percent read content in individual samples. Changing the order of alignment to reference genome does not change the results significantly

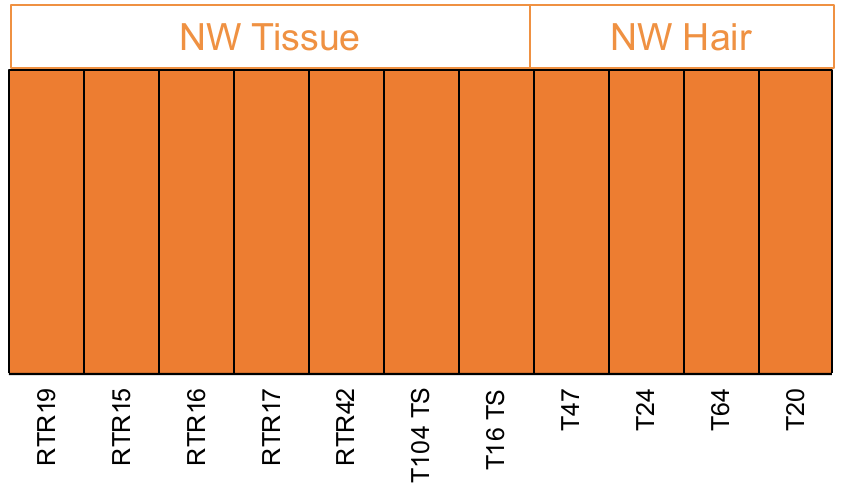

**Supplementary figure 3:** To test if biases were subtle enough to be detected at an intra-population scale structure was estimated using the shed hair and the Ranthambore samples in Natesh et al (2017). However, no structure is detected thus there are biases to create population level errors.
